# Quantifying Adipose Tissue Thermogenesis Using Highly Sensitive Isothermal Microcalorimetry

**DOI:** 10.64898/2026.01.07.698169

**Authors:** Pauke C. Schots, Devesh Kesharwani, Chad C. Doucette, Aaron C. Brown

**Author notes:** Corresponding author: Aaron C. Brown. These authors contributed equally to this work (Pauke C. Schots, Devesh Kesharwani). **Author Contributions:** P.C.S., D.K., and A.C.B. designed research; P.C.S, D.K. and C.C.D. performed research; A.C.B. contributed new reagents/analytic tools; P.C.S, D.K., C.C.D. and A.C.B. analysed data; A.C.B. P.C.S. and D.K. wrote the paper. **Competing Interest Statement:** The authors declare no competing interest.

## Abstract

Isothermal microcalorimetry enables direct measurement of thermogenic heat production, but optimal conditions for its application to adipose tissue have not been systematically defined. Here, we establish experimental parameters for quantifying thermogenesis using the CalScreener, a high-throughput isothermal microcalorimetry platform, across adipocyte organoids, freshly isolated adipocytes, and intact adipose tissue explants. Heat output scaled with spheroid size within a defined range and increased linearly with spheroid number per well, improving measurement consistency. Freshly isolated adipocytes and intact adipose tissue exhibited robust and depot-specific thermogenic activity that was sensitive to physiological modulation. Notably, intact adipose tissue retained functional thermogenic capacity for at least 6 hours *ex vivo* when maintained in nutrient-containing medium, enabling batch analysis of large cohorts. In contrast, storage conditions designed to preserve mitochondrial integrity during cold handling suppressed basal heat production and did not maintain integrated thermogenic activity, despite preserved adrenergic responsiveness. Together, these results provide practical guidance for tissue handling, assay design, and interpretation of isothermal microcalorimetry-based measurements of adipose tissue thermogenesis.

## Introduction

Obesity is a major global health challenge that increases the risk of metabolic disorders including type 2 diabetes, cardiovascular disease, metabolic-associated steatotic liver disease and cancer^1-3^. At a physiological level, obesity reflects a chronic imbalance between energy intake and energy expenditure that is driven in part by dysfunction of adipose tissue, a central regulator of systemic metabolic homeostasis^4^.

While lifestyle modification remains the cornerstone of obesity management, long-term efficacy is limited. Although pharmacological and surgical interventions can be effective, they are costly, associated with adverse effects, and frequently followed by weight regain^5^. These limitations have motivated increased interest in strategies that enhance energy expenditure rather than restricting energy intake.

Thermogenic adipocytes, including brown adipocytes and inducible beige adipocytes within white adipose tissue, dissipate chemical energy as heat and play a critical role in metabolic regulation^6-8^. Brown adipocytes reside in dedicated depots and are specialized for constitutive thermogenesis, whereas beige adipocytes emerge within subcutaneous white adipose tissue in response to environmental cues such as cold exposure^7^. Thermogenic activation is driven largely by sympathetic signaling through beta-adrenergic receptors, leading to increased substrate uptake, mitochondrial activity, and heat production^9, 10^. Although uncoupling protein 1 (UCP1) is a central mediator of thermogenesis, additional UCP1-independent mechanisms, including calcium cycling, creatine turnover, and futile lipid cycling, also contribute to energy dissipation^11^.

Despite the recognized importance of thermogenic adipocytes, accurately quantifying thermogenic activity remains technically challenging. Traditional approaches rely on surrogate measures such as gene and protein expression, oxygen consumption assays using Seahorse or Oroboros platforms, or glucose uptake assessed by fluorine-18 fluorodeoxyglucose positron emission tomography (PET) imaging^12-14^. While informative, these methods do not directly measure heat production, which is the defining functional output of thermogenesis. Recent advances in isothermal microcalorimetry have enabled direct, label-free measurement of heat production in living cells and tissues, providing an integrated readout of metabolic activity. Prior studies have demonstrated the feasibility of using the CalScreener isothermal microcalorimeter to quantify heat production in isolated primary and cell line-derived adipocytes, highlighting the potential of this approach for studying adipose tissue metabolism^15, 16^. However, key experimental parameters governing the application of isothermal microcalorimetry to adipose biology remain poorly defined. These include optimal cell and tissue configurations, the influence of three-dimensional organization, the stability of thermogenic activity in intact tissues *ex vivo*, and the impact of tissue handling and storage conditions on calorimetric readouts. In particular, it is not well established how long intact adipose tissue can retain functional thermogenic capacity outside the organism, which is a practical consideration for studies requiring extended tissue harvest times or batch analysis of large experimental cohorts.

In this study, we systematically evaluate the use of isothermal microcalorimetry to measure thermogenesis across multiple adipose tissue models, including three-dimensional adipocyte organoids, freshly isolated adipocytes, and intact adipose tissue explants from distinct depots. We define key parameters influencing heat production, assess depot-specific thermogenic behaviour, and examine how tissue storage conditions affect calorimetric measurements. Together, this work provides a practical methodological framework for the application of isothermal microcalorimetry to studies of adipose tissue thermogenesis and metabolic function.

## Results

### Optimization of adipocyte organoid size and number for calorimetric measurement of thermogenesis

We first assessed the sensitivity of the CalScreener in detecting heat production from three-dimensional adipocyte organoids, which offer the advantage of precise control over total cell number compared to intact adipose tissue explants. Brown adipocyte organoids were generated using cell attachment-free wells by seeding between 5,000 and 50,000 mouse preadipocytes from an immortalized ThermoMouse-derived cell line per organoid, followed by differentiation into mature brown adipocytes^17^. Individual organoids were seeded into CalWell vessels, with three technical replicates per organoid size. Heat production increased in a dose-dependent manner as the number of cells seeded per organoid increased from 5,000 to 30,000 cells, plateaued between 30,000 and 40,000 cells, and declined at 50,000 cells (Fig. 1A-B). A similar size-dependent relationship was observed using an immortalized beige preadipocyte cell line after differentiation into mature adipocytes (Fig. 1C-D).

**Figure 1.**
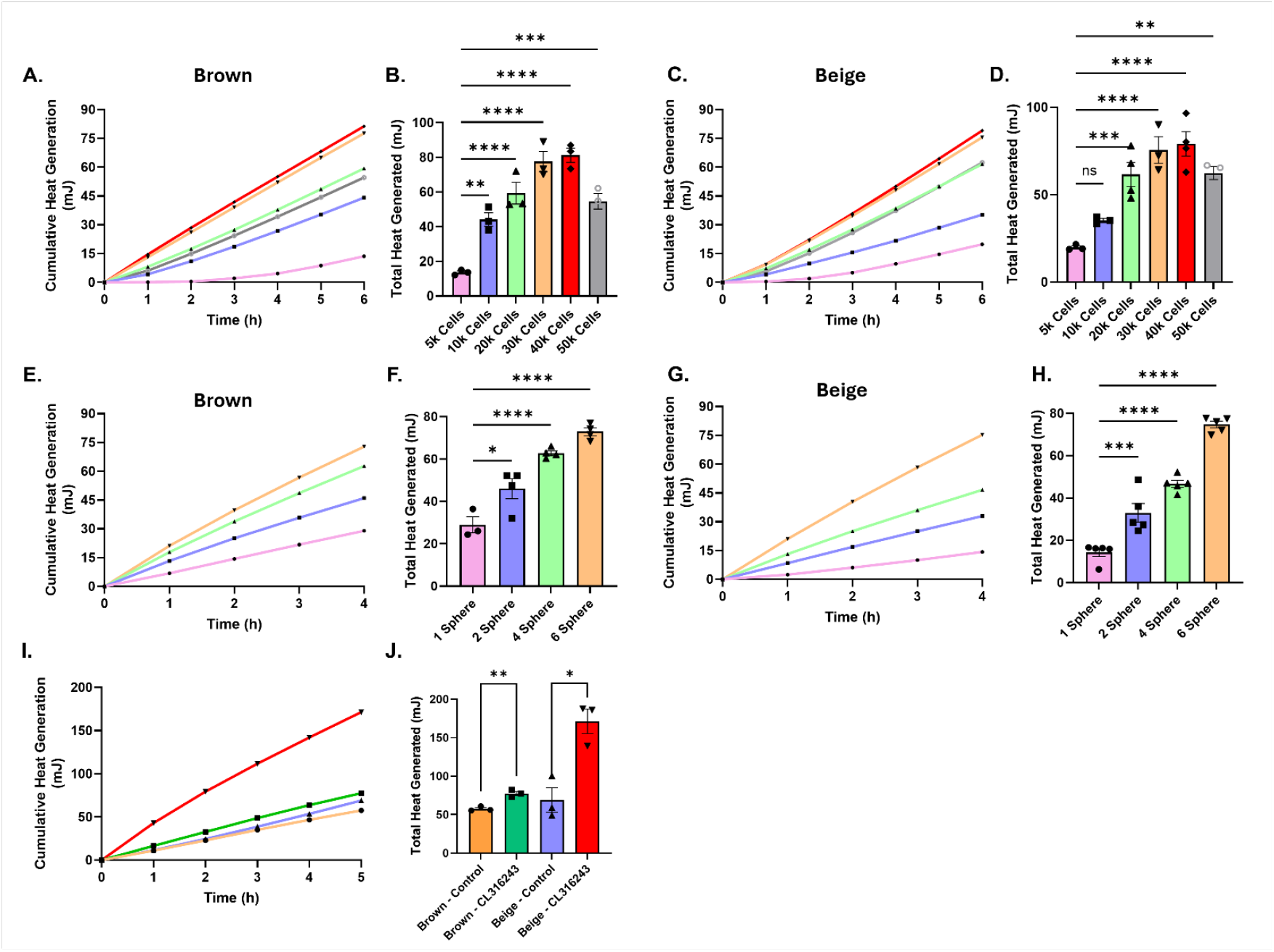
Optimization of adipocyte organoid size and number for calorimetric measurement of thermogenesis. (A) Cumulative heat production over time from differentiated brown adipocyte organoids generated from 5,000–50,000 preadipocytes per organoid. (B) Total heat production from brown adipocyte organoids of increasing size. (C) Cumulative heat production over time from differentiated beige adipocyte organoids generated across the same range of initial cell numbers. (D) Total heat production from beige adipocyte organoids of increasing size. (E) Cumulative heat production over time from brown adipocyte organoids with increasing numbers of organoids (1–6) per CalWell. (F) Total heat production from brown adipocyte organoids as a function of organoid number per well. (G) Cumulative heat production over time from beige adipocyte organoids with increasing numbers of organoids per CalWell. (H) Total heat production from beige adipocyte organoids as a function of organoid number per well. (I) Cumulative heat production over time from pooled brown and beige adipocyte organoids treated with vehicle or the β3-adrenergic agonist CL316243. (J) Total heat production from brown and beige adipocyte organoids following vehicle or CL316243 treatment. Data are shown as mean ± SEM. Individual data points represent independent CalWell measurements. Statistical analyses were performed using Student’s t-test and one-way ANOVA as indicated.

Although we did not directly assess viability, oxygen gradients, or necrosis within organoids in these experiments, prior studies in three-dimensional organoid systems have demonstrated that increasing spheroid size can lead to diffusion limitations for oxygen and nutrients, resulting in reduced cellular function or central necrosis in larger structures^18, 19^. An alternative explanation for the reduced heat signal at the highest cell number could be depletion of nutrients or oxygen within the CalWell microenvironment.

However, across the 6-hour measurement period, heat production increased approximately linearly over time for all organoid sizes, including those seeded with up to 50,000 cells, suggesting that gross medium depletion was unlikely to be limiting under our experimental conditions (Fig. 1A-D).

Based on these findings, we next tested whether adding multiple organoids per CalWell could reduce variability and improve measurement robustness. Organoids seeded at 30,000 preadipocytes per spheroid were differentiated into either brown or beige adipocytes, and between one and six organoids were added per CalWell (Fig. 1E-H). Heat production increased linearly with the number of organoids per well for both brown and beige adipocyte organoids. Notably, wells containing four to six organoids exhibited reduced variability across replicate wells compared with wells containing one or two organoids. Pooled organoids also responded to β3-adrenergic stimulation, as evidenced by increased heat production (Fig. 1I-J).

Culture-derived brown and beige adipocyte organoids from our chosen cell lines exhibited similar basal heat output; however, beige adipocytes responded more robustly to CL316243. This likely reflects properties of the individual clonal cell lines rather than a generalizable difference between brown and beige adipocytes.

Together, these data demonstrate that the CalScreener is sufficiently sensitive to detect heat production from thermogenic adipocytes organized in a three-dimensional architecture across a defined range of cell numbers, that excessive organoid size is associated with a plateau or decline in heat output, and that inclusion of multiple organoids per well improves measurement consistency. These findings establish key experimental parameters for the use of adipocyte organoids in calorimetric assays of thermogenic function.

### Sustained thermogenic responsiveness of freshly isolated adipocytes measured by isothermal microcalorimetry

We next moved to *ex vivo* studies to determine whether heat production from adipocytes freshly isolated from mouse adipose tissue could be directly measured and whether these cells retained responsiveness to a thermogenic agonist. Interscapular brown (iBAT) and subcutaneous inguinal adipose tissue (iWAT) depots were pooled from multiple mice and enzymatically digested to release mature adipocytes. The floating adipocyte fraction was collected, and equivalent numbers of adipocytes (based on counting) were loaded per well for microcalorimetric analysis using the CalScreener. Mature brown adipocytes exhibited a 140% increase in heat production in response to the β3-adrenergic agonist CL316243, with thermogenesis continuing to rise through the end of the 6-hour CalScreener run (Fig. 2). Mature inguinal-derived adipocytes showed a more modest but significant ∼50% increase in heat production that also continued to increase over the 6-hour measurement period (Fig. 2). Together, these results demonstrate that adipocytes freshly isolated from mouse adipose tissue remain viable and capable of mounting sustained thermogenic responses *ex vivo* for an extended period of time, encompassing tissue harvest, treatment and loading, and microcalorimetric measurement.

**Figure 2.**
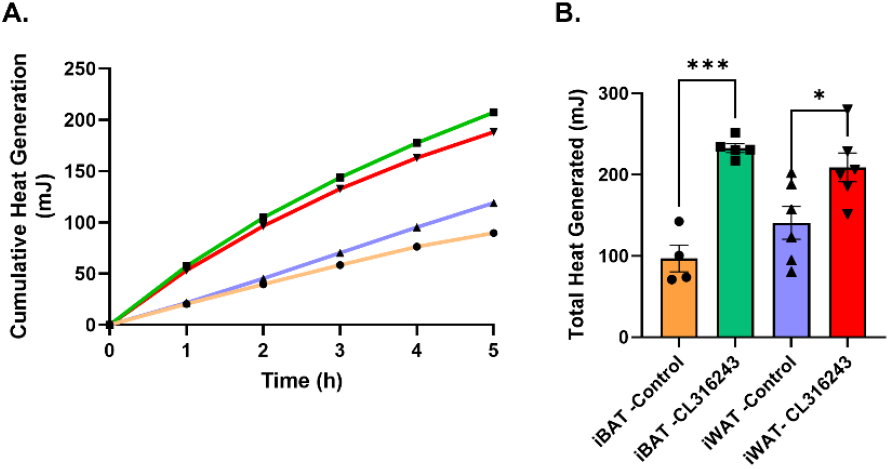
Sustained thermogenic responsiveness of freshly isolated adipocytes measured by isothermal microcalorimetry. (A) Cumulative heat production over time from mature adipocytes freshly isolated from interscapular brown adipose tissue (iBAT) or subcutaneous inguinal adipose tissue (iWAT) and treated with vehicle or the β3-adrenergic agonist CL316243. (B) Total heat production from freshly isolated brown and inguinal adipocytes following vehicle or CL316243 treatment. Data are shown as mean ± SEM. Individual data points represent independent CalWell measurements. Statistical significance was assessed by one-way ANOVA as indicated. Abbreviations: iBAT, interscapular brown adipose tissue; iWAT, inguinal white adipose tissue; gWAT, gonadal white adipose tissue.

### Depot-specific thermogenic profiles of intact adipose tissue explants and modulation by thermoneutral housing

We next tested freshly isolated intact adipose tissue explants from mice housed at room temperature using microcalorimetry over a 16-hour period. Adipose depots were rapidly harvested and immediately placed in DMEM prior to analysis. Freshly isolated iBAT exhibited an approximately 350% increase in heat output per gram of tissue compared with either iWAT or gonadal adipose tissue (gWAT), which generated similar levels of heat under these conditions (Fig. 3). When mice were housed at thermoneutrality for 7 days prior to tissue harvest, heat production was significantly reduced across all adipose depots, although the overall relative pattern was preserved (BAT > inguinal ≈ gonadal) (Fig. 3). Brown adipose tissue exhibited the most pronounced reduction, losing approximately 70% of its heat production relative to room temperature housed mice, whereas iWAT and gWAT each lost approximately 50%. This differential reduction is consistent with the greater basal thermogenic capacity of iBAT, which is more strongly suppressed at thermoneutrality, while browning within WAT depots is relatively limited even at room temperature.

**Figure 3.**
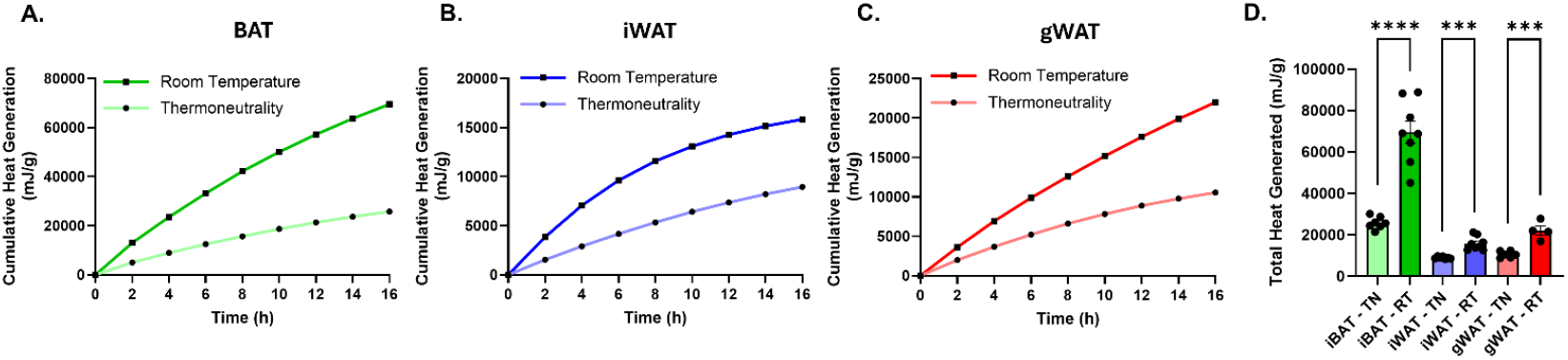
Depot-specific thermogenic activity of intact adipose tissue explants and suppression by thermoneutral housing. (A-C) Cumulative heat production over time, normalized per gram of tissue, measured by isothermal microcalorimetry from freshly isolated intact adipose tissue explants of iBAT (A), iWAT (B), gWAT (C) harvested from mice housed at room temperature or thermoneutrality. (D) Total heat production per gram of tissue from iBAT, iWAT, and gWAT explants under each housing condition. Data are shown as mean ± SEM. Individual data points represent independent tissue explants. Statistical significance was assessed by Student’s t-test as indicated.

Unexpectedly, gWAT consistently exhibited heat production that was equivalent to or slightly higher than that of iWAT, despite the well-established association of subcutaneous depots with greater browning potential in response to cold exposure^20, 21^. Because iWAT is known to exhibit higher thermogenic capacity *in vivo* at room temperature compared with gWAT, these findings are unlikely to reflect intrinsically greater thermogenesis within the gonadal depot. Instead, a more plausible explanation is that gWAT may be more resistant to progressive loss of metabolic activity under sealed microcalorimetry conditions. Consistent with this interpretation, iWAT and gWAT displayed similar initial rates of heat generation early in the time course; however, heat accumulation in iWAT plateaued over the 16-hour measurement period, whereas gWAT maintained a more sustained increase in heat output. Together, these observations indicate that depot-specific differences in tissue longevity or stress resistance, rather than differences in intrinsic thermogenic programming, may contribute to the relative heat production observed during extended *ex vivo* microcalorimetry.

### Assessment of regional thermogenic uniformity within inguinal adipose tissue

The iWAT depot has been reported to exhibit spatial heterogeneity in thermogenic potential and *Ucp1* expression along its longitudinal axis during browning^22^. To determine whether regional differences in thermogenic capacity could be detected *ex vivo*, we sectioned iWAT depots longitudinally and transversely to generate anatomically distinct subsections (Fig. 4A). In mice housed at room temperature, no significant differences in heat production were observed between left versus right longitudinal sections or between upper versus lower transverse sections of the inguinal depot (Fig. 4B-C). These data suggest that under basal room temperature conditions, thermogenic capacity is relatively uniform across the inguinal adipose tissue depot.

**Figure 4.**
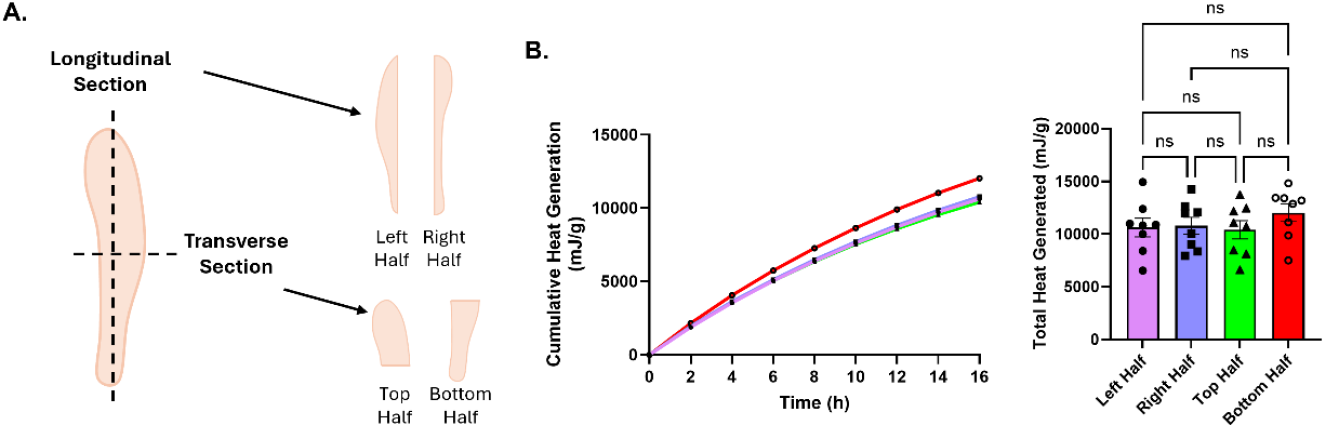
Assessment of regional thermogenic uniformity within inguinal adipose tissue. (A) Schematic illustrating longitudinal and transverse sectioning of the iWAT) depot to generate anatomically distinct subsections. (B) Cumulative heat production over time, normalized per gram of tissue, measured by isothermal microcalorimetry from ex vivo iWAT subsections corresponding to left and right longitudinal halves and upper and lower transverse halves. (C) Total heat production per gram of tissue from each iWAT subsection. Data are shown as mean ± SEM. Individual data points represent independent tissue sections. Statistical significance was assessed by one-way ANOVA as indicated.

### Effects of tissue storage conditions on basal thermogenesis and adrenergic responsiveness in intact adipose explants

Many metabolic studies require the harvest and analysis of adipose tissues from large cohorts of mice, which can necessitate extended tissue harvest times. To assess whether short-term storage of adipose tissue prior to isothermal microcalorimetry affects relative heat production, freshly harvested tissues were stored for up to 6 hours in either nutrient-containing DMEM at 37C or on ice, and compared with freshly isolated tissue placed directly into the CalScreener. All tissues were loaded simultaneously to enable direct comparison of heat production. Storage in DMEM at 37C did not significantly impair thermogenic capacity in iBAT or iWAT relative to freshly isolated controls, whereas storage on ice significantly reduced heat production (Fig. 5A-D). gWAT exhibited greater variability but showed a similar trend toward reduced heat output following cold storage (Fig. 5E-F). Because BIOPS (Biopsy Preservation Solution) buffer preserves mitochondrial integrity and respiratory capacity during cold handling, we tested whether cold storage of mitochondria-rich iBAT in BIOPS preserves thermogenic capacity. Contrary to our hypothesis, tissue stored in BIOPS on ice for 6 hours exhibited reduced heat production compared with tissue maintained in DMEM, despite all samples being assayed in DMEM during the CalScreener run (Fig. 5G-H). These findings indicate that preservation of mitochondrial integrity and respiratory competence during cold storage is not sufficient to maintain integrated thermogenic activity in intact brown adipose tissue explants.

**Figure 5.**
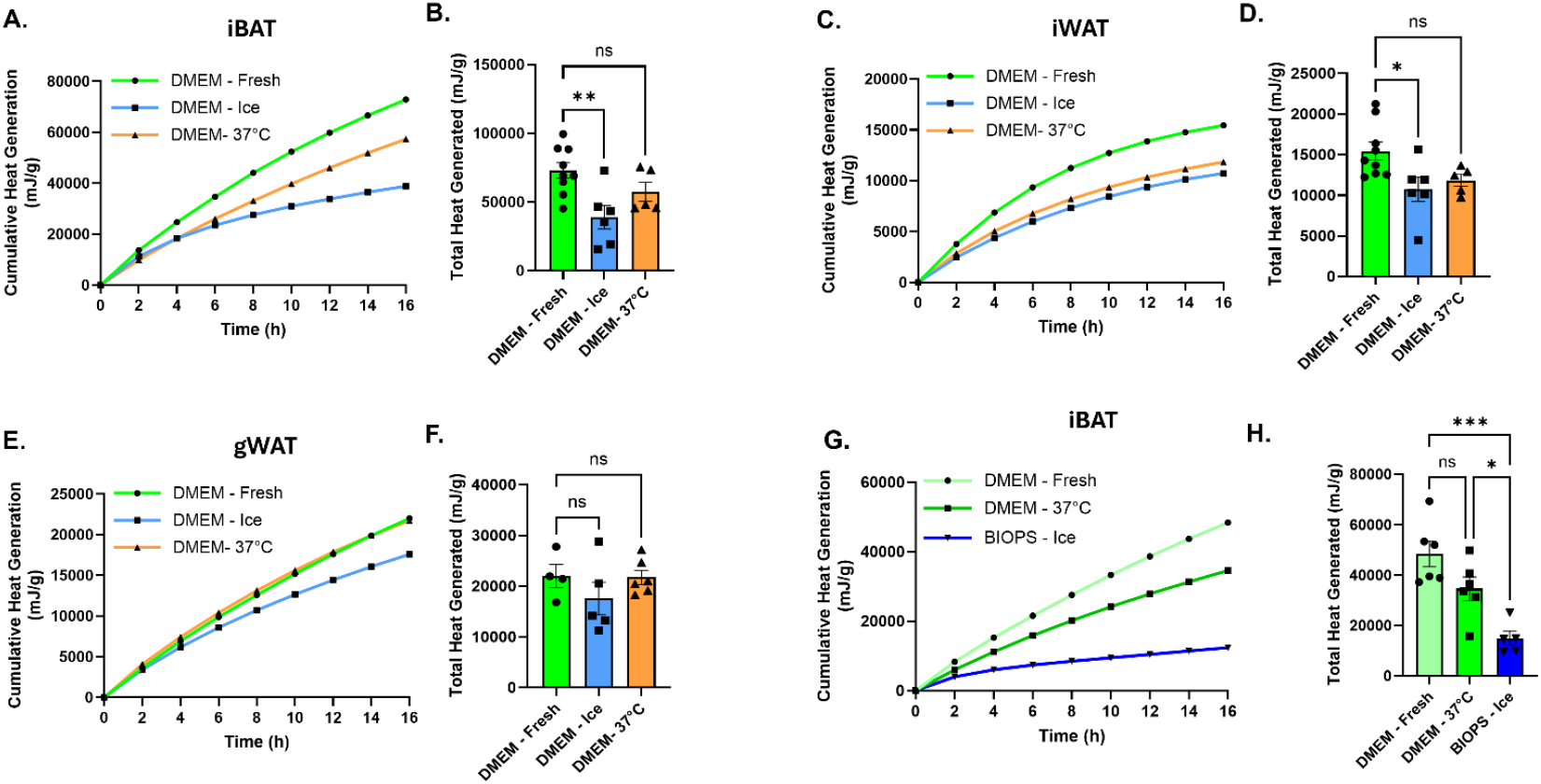
Effects of tissue storage conditions on basal thermogenesis in intact adipose tissue explants. (A,B) Cumulative heat production over time (A) and total heat production (B), normalized per gram of tissue, from iBAT explants that were freshly isolated or stored for 6 hours in DMEM on ice or at 37C prior to isothermal microcalorimetry. (C,D) Cumulative heat production (C) and total heat production (D), normalized per gram of tissue, from iWAT explants under the same storage conditions. (E,F) Cumulative heat production (E) and total heat production (F), normalized per gram of tissue, from gWAT explants following fresh isolation or short-term storage in DMEM on ice or at 37 °C. (G,H) Cumulative heat production (G) and total heat production (H), normalized per gram of tissue, from iBAT explants that were freshly isolated or stored for 6 hours in either DMEM at 37 °C or BIOPS buffer (fresh or on ice) prior to analysis; all samples were assayed in DMEM during the CalScreener run. Data are shown as mean ± SEM. Individual data points represent independent tissue explants. Statistical significance was assessed by one-way ANOVA as indicated.

Finally, we tested the ability of iBAT explants to respond to the pan β-adrenergic agonist isoproterenol under three conditions: freshly isolated iBAT, iBAT stored for 6 hours in DMEM at 37C, and iBAT stored for 6 hours in BIOPS on ice (Fig. 6). Basal heat production was approximately two-fold higher in fresh tissue and in iBAT stored in DMEM at 37C compared with BAT stored in BIOPS on ice (Fig. 6B-D). Despite these baseline differences, isoproterenol significantly increased heat production in all three conditions. Fresh iBAT and iBAT stored in DMEM at 37C for 6 hours exhibited comparable increases in heat production of approximately 2-fold relative to baseline, whereas iBAT stored in BIOPS on ice for 6 hours displayed a larger relative increase of approximately three-fold, restoring absolute heat production to levels comparable to those observed in isoproterenol-treated BAT stored in DMEM. Together, these data indicate that although cold storage in BIOPS suppresses basal thermogenic activity, it preserves β-adrenergic responsiveness of iBAT explants.

**Figure 6.**
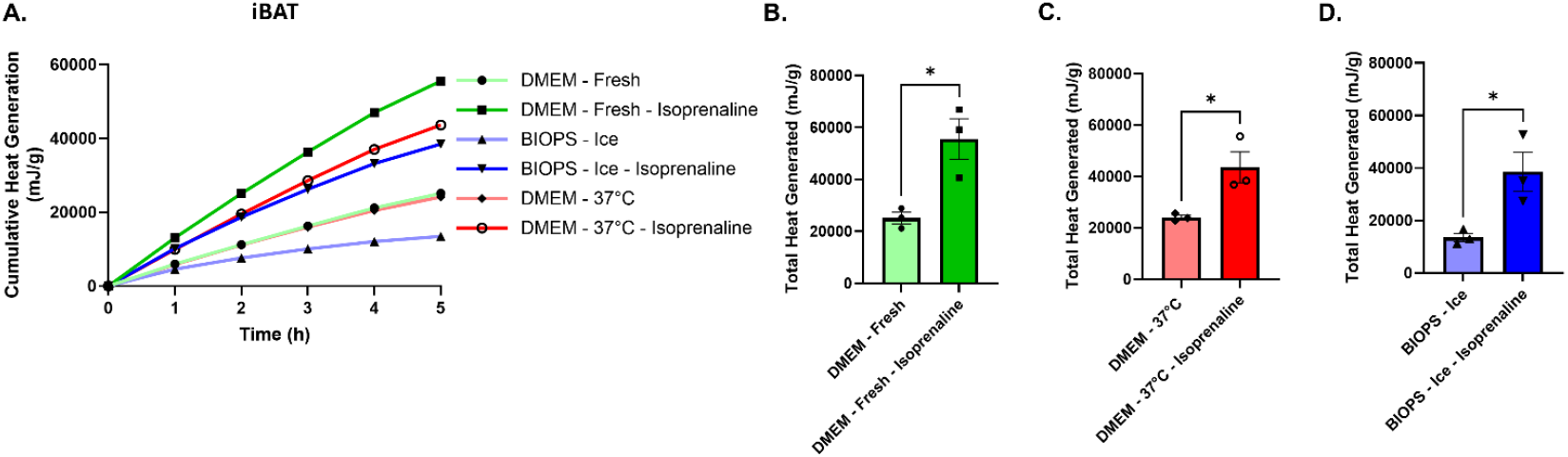
Preservation of β-adrenergic responsiveness following short-term storage of interscapular brown adipose tissue. (A) Cumulative heat production over time, normalized per gram of tissue, from iBAT explants that were freshly isolated or stored for 6 hours in DMEM at 37 °C or in BIOPS buffer on ice, and subsequently treated with vehicle or the pan β-adrenergic agonist isoproterenol. (B–D) Total heat production per gram of tissue from iBAT explants treated with vehicle or isoproterenol following fresh isolation (B), storage in DMEM at 37 °C for 6 hours (C), or storage in BIOPS on ice for 6 hours (D). Data are shown as mean ± SEM. Individual data points represent independent tissue explants. Statistical significance was assessed by Student’s t-test as indicated.

## Discussion

In this study, we establish isothermal microcalorimetry using the CalScreener as a robust platform for quantifying thermogenic heat production across a spectrum of adipose tissue models, ranging from three-dimensional adipocyte organoids to freshly isolated adipocytes and intact adipose tissue explants.

Together, these data define key experimental parameters, clarify technical limitations, and provide practical guidance for the application of microcalorimetry to studies of adipose tissue thermogenesis. Using three-dimensional brown and beige adipocyte organoids, we demonstrate that calorimetric heat output scales predictably with cell number across a defined range, with a plateau and decline at larger organoid sizes.

This behaviour is consistent with diffusion constraints and functional limitations previously described in spheroid systems and underscores the importance of optimizing organoid size for metabolic assays^18, 19^. Importantly, pooling multiple organoids per well reduced variability and improved assay robustness, supporting the use of multi-organoid loading strategies for reproducible calorimetric measurements. These findings establish adipocyte organoids as a tractable and scalable system for calorimetric interrogation of thermogenic function.

Extending these approaches to *ex vivo* preparations, we show that mature adipocytes freshly isolated from brown and subcutaneous adipose tissue remain viable and thermogenically responsive for extended periods *ex vivo*, mounting sustained heat production in response to β3-adrenergic stimulation. Notably, intact adipose tissue explants retained measurable and physiologically meaningful thermogenic activity for at least 6 hours *ex vivo* when maintained in nutrient-containing medium at physiological temperature. To our knowledge, the temporal durability of functional thermogenesis in intact adipose tissue has not been systematically evaluated, and these results establish that thermogenic capacity persists over experimentally relevant timeframes compatible with large-cohort tissue harvests and batch microcalorimetric analysis.

Consistent with known *in vivo* physiology, intact adipose tissue explants exhibited depot-specific differences in thermogenic output, with brown adipose tissue displaying the highest basal heat production and thermoneutral housing markedly suppressing thermogenesis across depots. Brown adipose tissue showed the greatest reduction following thermoneutral housing, reflecting its higher dependence on sympathetic activation, whereas white adipose depots exhibited more modest changes. These findings indicate that isothermal microcalorimetry captures physiologically relevant modulation of thermogenic state in intact tissues *ex vivo*. Prolonged microcalorimetric measurements further revealed depot-specific differences in the stability of metabolic activity under sealed assay conditions. Despite the greater browning potential of inguinal adipose tissue *in vivo*, gonadal adipose tissue-maintained heat production more effectively over extended time courses. This suggests that differences in tissue resilience to nutrient depletion, oxygen availability, or metabolic stress may influence long-duration calorimetric readouts and highlights an important consideration when interpreting absolute heat production across adipose depots *ex vivo*.

A key practical outcome of this work is the systematic evaluation of tissue storage conditions prior to isothermal microcalorimetry. We demonstrate that short-term storage of adipose tissue for up to 6 hours in nutrient-containing DMEM at physiological temperature preserves thermogenic capacity comparably to freshly isolated tissue, enabling synchronized analysis of large experimental cohorts. In contrast, cold storage on ice markedly suppresses basal heat production. BIOPS is a physiological buffer widely used to preserve mitochondrial structure and respiratory capacity during *ex vivo* handling, owing to its defined ionic composition, inclusion of ATP and phosphocreatine, and buffering of calcium, which together stabilize mitochondrial integrity during short-term cold storage^23, 24^. However, BIOPS lacks exogenous metabolic substrates and is optimized for maintaining mitochondrial competence rather than supporting active cellular metabolism. Importantly, following storage, all tissues, including those stored in BIOPS, were returned to nutrient-containing DMEM for isothermal microcalorimetry. Under these equivalent assay conditions, BIOPS-stored tissues nonetheless exhibited reduced basal thermogenic activity, indicating that preservation of mitochondrial integrity during cold storage does not maintain integrated thermogenic capacity in intact adipose tissue explants.

Importantly, despite reduced basal heat output following BIOPS cold storage, β-adrenergic responsiveness was preserved, with explants exhibiting robust stimulation of heat production upon isoproterenol treatment. This suggests that cold storage in BIOPS primarily suppresses basal metabolic activity rather than irreversibly impairing thermogenic signaling pathways or mitochondrial capacity. Conceptually, these findings reinforce the idea that thermogenesis measured by isothermal microcalorimetry reflects a coordinated, systems-level metabolic output that depends on intact cellular signaling, substrate availability, and mitochondrial function, rather than mitochondrial integrity alone.

### Limitations and Future Directions

Several limitations of this study should be acknowledged. First, while isothermal microcalorimetry provides a sensitive and integrative measure of heat production, it does not directly resolve the underlying metabolic pathways contributing to thermogenesis. Future studies combining microcalorimetry with molecular, biochemical, and imaging-based approaches will be important for linking heat output to specific thermogenic mechanisms, such as UCP1-dependent versus UCP1-independent pathways. Second, the sealed nature of the CalScreener wells may impose constraints on oxygen and nutrient availability during prolonged measurements, which could differentially affect adipose depots and contribute to the depot-specific divergence observed over extended time courses. Incorporation of defined substrate supplementation or limiting microcalorimetry measurements to shorter time windows may help reduce nutrient depletion and metabolic stress, and thereby further clarify depot-specific differences in thermogenic stability observed during prolonged assays. Third, although we demonstrate preservation of thermogenic capacity for up to 6 hours *ex vivo*, longer storage durations were not examined. Defining the upper temporal limits of functional thermogenesis, as well as identifying conditions that extend tissue viability without compromising metabolic activity, represents an important area for future methodological development. Finally, the experiments presented here focused primarily on murine adipose tissue and immortalized cell lines. Extension of these approaches to primary human adipose tissue, human adipocyte organoids, and disease-relevant models will be critical for translating isothermal microcalorimetry into broader metabolic and translational research applications.

## Conclusions

Overall, this work establishes isothermal microcalorimetry as a sensitive and versatile approach for studying adipose tissue thermogenesis across multiple experimental models while defining key technical parameters related to sample configuration, assay duration, and tissue handling conditions. By demonstrating that intact adipose tissue retains functional thermogenic capacity over extended *ex vivo* timeframes, these findings provide an important methodological framework for future studies of adipose tissue metabolism and its regulation under physiological and pathological conditions.

## Acknowledgements

This work was supported by National Institute of Diabetes and Digestive and Kidney Diseases (NIDDK) award R01DK124261 (A. Brown) and by the SECURE project from UiT—The Arctic University of Norway, grant ID 2061344. This research utilized equipment supported by NIH award U54GM115516 (C. J. Rosen, Principal Investigator).

## Methods

### Reagents and Buffers

BIOPS (biopsy preservation solution) buffer was prepared as a physiological preservation buffer containing 10 mM Ca-EGTA buffer (0.1 µM free Ca^2+^), 20 mM imidazole, 20 mM taurine, 50 mM K-MES, 0.5 mM dithiothreitol (DTT), 6.56 mM MgCl_2_, 5.77 mM ATP, and 15 mM phosphocreatine, adjusted to pH 7.1. Dulbecco’s modified Eagle’s medium (DMEM; high glucose) supplemented with GlutaMAX™ and HEPES (Gibco™, 32430027) and 1% bovine serum albumin (BSA; fraction V) was used for tissue incubation and calorimetric measurements. CL316243 was purchased from Cayman Chemicals (17499). Isoproterenol hydrochloride was purchased from Sigma-Aldrich (I5627).

### Animal Ethics and Husbandry

All animal studies were conducted in accordance with applicable national and institutional guidelines and were approved by the relevant animal ethics committees at each study site. Experiments performed in the United States were conducted in compliance with the National Institutes of Health Guide for the Care and Use of Laboratory Animals and were approved by the Institutional Animal Care and Use Committee (IACUC; protocol #2203) at the MaineHealth Institute for Research (MHIR).

Experiments performed in Norway were approved by the institutional animal use authority at the Faculty of Health Sciences, UiT - The Arctic University of Norway (approval reference AKM 04/25), and were conducted in accordance with Directive 2010/63/EU on the protection of animals used for scientific purposes. All mice used in this study were adult C57BL/6 animals aged 8-11 weeks. Mice were housed in pathogen-free facilities under controlled environmental conditions with a 12-hour light/dark cycle and ad libitum access to food and water. Animals euthanized at MHIR were sacrificed by carbon dioxide (CO_2_) inhalation in accordance with IACUC-approved protocols, whereas animals euthanized at UiT were sacrificed by overdose of pentobarbital. All efforts were made to minimize animal suffering and to reduce animal numbers wherever possible.

### *In Vitro* Adipocyte Culture and Differentiation

Three-dimensional adipocyte organoids were generated using attachment-free spheroid culture plates (Greiner Bio-One CELLSTAR 96-well microplate, 650979). Immortalized interscapular brown (ThermoMouse) and mouse inguinal adipose tissue derived beige (Kerafast, Inc. Cat# EVC005) preadipocyte cell lines were seeded at defined cell numbers per well and allowed to self-assemble into organoids prior to differentiation^17, 25^. Cells were then induced to differentiate using a defined adipogenic cocktail containing insulin, triiodothyronine (T3), rosiglitazone, IBMX, dexamethasone, indomethacin, SB431542, and ascorbic acid phosphate as previously described^16, 26^. After 3 days, cultures were transitioned to maintenance medium containing insulin, T3, rosiglitazone, SB431542, and ascorbic acid phosphate for 3 days, and were maintained until day 10 of differentiation in basal medium (DMEM + 10% FBS) prior to CalScreener analysis.

Mature adipocytes were isolated from interscapular brown, inguinal white and gonadal white adipose tissue of adult mice. Dissected adipose tissues were minced and digested at 37 °C for 30-40 minutes in tissue lysis buffer containing 0.123 M NaCl, 1.3 mM CaCl_2_, 5 mM glucose, 100 mM HEPES, 4% (v/v) bovine serum albumin (BSA; fraction V), and 0.1% (w/v) Collagenase P with constant rocking. Following digestion, an equal volume of autoMACS® Running Buffer was added to each sample to inhibit the collagenase activity. Tissue lysates were then allowed to stand at room temperature for 10 minutes to facilitate separation of mature adipocytes, which floated to the upper fraction. Mature adipocytes were carefully collected from the floating layer using a 22-gauge syringe and washed with sterile autoMACS® Running Buffer to remove residual debris and stromal cells. Cell viability and concentration were determined using the Cellometer K2 Fluorescent Cell Counter (Revity, Inc) following staining with ViaStain™ AOPI solution, according to the manufacturer’s instructions.

### Adipose Tissue Dissection and Preparation for *Ex Vivo* Tissue Analysis

Interscapular BAT, inguinal WAT, and gonadal WAT were rapidly dissected, cleaned of visible connective tissue, and sectioned using sterile scalpel blades. Tissue sections were weighed prior to incubation in the Calscreener. Typical BAT sections ranged from approximately 4 to 22 mg, whereas WAT sections ranged from approximately 40 to 175 mg depending on depot. Tissue sections were immediately placed into DMEM or BIOPS buffer depending on the experimental condition.

### Temporary Storage of Adipose Tissue

To evaluate the effects of short-term storage prior to calorimetric analysis, adipose tissue sections were stored for up to 6 hours under defined conditions. Storage conditions included incubation in nutrient-containing DMEM at 37C or cold storage on ice in either DMEM or BIOPS buffer. Tissues assigned to storage conditions were harvested at the beginning of the storage period, whereas freshly isolated control tissues were harvested near the end of the same time window. All tissues were loaded into the CalScreener simultaneously to allow direct comparison. For calorimetric measurements, all tissues, including those previously stored in BIOPS, were transferred into DMEM prior to loading, ensuring equivalent nutrient availability during the microcalorimetry run.

### Isothermal Microcalorimetry

Heat production was measured using a 48 well isothermal microcalorimeter (calScreener™, Symcel, Sweden). Plastic inserts were filled with 200 µL DMEM, adjusted to account for tissue volume by assuming a tissue density of 1 mg/µL to maintain comparable oxygen availability across samples. Tissue sections, isolated adipocytes, or adipocyte organoids were placed into the inserts, which were then sealed within titanium calVial™ capsules and loaded into the calPlate™ holder. When indicated, CL316243 (2 µM) or isoproterenol (500 µM) was added immediately prior to sealing the vials. After insertion into the instrument, samples underwent a 30-minute equilibration and calibration period, followed by an additional 15-minute stabilization period before continuous heat flow was recorded. Measurements were performed for 6-16 hours depending on the experimental condition.

### Statistical Analysis

All data are presented ± standard error of the mean (SEM). Comparisons between two groups were performed using unpaired two-tailed Student’s t-tests. For comparisons involving more than two groups, one-way analysis of variance (ANOVA) was used followed by Tukey’s multiple-comparison test where appropriate. A p-value < 0.05 was considered statistically significant. Significance levels are indicated as follows: p < 0.05 (*), p < 0.01 (**), p < 0.001 (***), and p < 0.0001 (****).

## References

1. Lumeng, C.N. & Saltiel, A.R. Inflammatory links between obesity and metabolic disease. J Clin Invest 121, 2111–2117 (2011).

2. Powell-Wiley, T.M. et al. Obesity and cardiovascular disease: a scientific statement from the American Heart Association. Circulation 143, e984–e1010 (2021).

3. Pati, S., Irfan, W., Jameel, A., Ahmed, S. & Shahid, R.K. Obesity and cancer: a current overview of epidemiology, pathogenesis, outcomes, and management. Cancers 15, 485 (2023).

4. Kajimura, S., Spiegelman, B.M. & Seale, P. Brown and beige fat: physiological roles beyond heat generation. Cell metabolism 22, 546–559 (2015).

5. Lauti, M., Kularatna, M., Hill, A.G. & MacCormick, A.D. Weight Regain Following Sleeve Gastrectomy-a Systematic Review. Obes Surg 26, 1326–1334 (2016).

6. Cypess, A.M. & Kahn, C.R. Brown fat as a therapy for obesity and diabetes. Curr Opin Endocrinol Diabetes Obes 17, 143–149 (2010).

7. Harms, M. & Seale, P. Brown and beige fat: development, function and therapeutic potential. Nat Med 19, 1252–1263 (2013).

8. Wu, J., Cohen, P. & Spiegelman, B.M. Adaptive thermogenesis in adipocytes: is beige the new brown? Genes Dev 27, 234–250 (2013).

9. Cero, C. et al. beta3-Adrenergic receptors regulate human brown/beige adipocyte lipolysis and thermogenesis. JCI Insight 6 (2021).

10. Rahbani, J.F. et al. ADRA1A-Galpha(q) signalling potentiates adipocyte thermogenesis through CKB and TNAP. Nat Metab 4, 1459–1473 (2022).

11. Cohen, P. & Kajimura, S. The cellular and functional complexity of thermogenic fat. Nat Rev Mol Cell Biol 22, 393–409 (2021).

12. Mahdaviani, K., Benador, I. & Shirihai, O. Assessment of Brown Adipocyte Thermogenic Function by High-throughput Respirometry. Bio Protoc 5 (2015).

13. Porter, C. et al. Human and Mouse Brown Adipose Tissue Mitochondria Have Comparable UCP1 Function. Cell Metab 24, 246–255 (2016).

14. Hankir, M.K. et al. Dissociation Between Brown Adipose Tissue (18)F-FDG Uptake and Thermogenesis in Uncoupling Protein 1-Deficient Mice. J Nucl Med 58, 1100–1103 (2017).

15. Bokhari, M.H. et al. Isothermal microcalorimetry measures UCP1-mediated thermogenesis in mature brite adipocytes. Commun Biol 4, 1108 (2021).

16. Doucette, C.C. et al. Optogenetic activation of UCP1-dependent thermogenesis in brown adipocytes. iScience 26, 106560 (2023).

17. Galmozzi, A. et al. ThermoMouse: an in vivo model to identify modulators of UCP1 expression in brown adipose tissue. Cell Rep 9, 1584–1593 (2014).

18. LaMontagne, E., Muotri, A.R. & Engler, A.J. Recent advancements and future requirements in vascularization of cortical organoids. Front Bioeng Biotechnol 10, 1048731 (2022).

19. Moss, S.P., Bakirci, E. & Feinberg, A.W. Engineering the 3D structure of organoids. Stem Cell Reports 20, 102379 (2025).

20. Vamvini, M. et al. Exercise training and cold exposure trigger distinct molecular adaptations to inguinal white adipose tissue. Cell Rep 43, 114481 (2024).

21. Paschos, G.K. et al. Cold-Induced Browning of Inguinal White Adipose Tissue Is Independent of Adipose Tissue Cyclooxygenase-2. Cell Rep 24, 809–814 (2018).

22. Wang, T. et al. Single-nucleus transcriptomics identifies separate classes of UCP1 and futile cycle adipocytes. Cell Metab 36, 2130–2145 e2137 (2024).

23. Canto, C. & Garcia-Roves, P.M. High-Resolution Respirometry for Mitochondrial Characterization of Ex Vivo Mouse Tissues. Curr Protoc Mouse Biol 5, 135–153 (2015).

24. Doerrier, C. et al. High-Resolution FluoRespirometry and OXPHOS Protocols for Human Cells, Permeabilized Fibers from Small Biopsies of Muscle, and Isolated Mitochondria. Methods Mol Biol 1782, 31–70 (2018).

25. Brown, A.C. Optogenetics Sheds Light on Brown and Beige Adipocytes. J Cell Signal 4, 178–186 (2023).

26. Su, S. et al. A Renewable Source of Human Beige Adipocytes for Development of Therapies to Treat Metabolic Syndrome. Cell Rep 25, 3215–3228 e3219 (2018).

